# Basolateral localization of Kcnj15 in branchial ionocytes of the seawater pufferfish *Takifugu rubripes*

**DOI:** 10.64898/2026.09.05.749310

**Authors:** Yohei Sakamoto, Akira Kato

**Affiliations:** School of Life Science and Technology, Institute of Science Tokyo, Yokohama-shi, Kanagawa 226-8501, Japan

**Keywords:** Kcnj15, ionocyte, potassium channel, osmoregulation, marine teleost

## Abstract

In marine teleosts, gill ionocytes play central roles in the excretion of excess monovalent ions (Na^+^, Cl^−^, and K^+^). Basolateral K^+^ channels are considered essential for sustaining ion secretion by maintaining K^+^ homeostasis and membrane potential; however, the molecular identity of basolateral K^+^ channels in teleost gill ionocytes remains incompletely understood. Kcnj15 is a K^+^ channel expressed in multiple osmoregulatory tissues of the Japanese pufferfish (*Takifugu rubripes*), and its localization changes depending on tissue type and osmotic conditions; however, its localization in the gills has remained unresolved. Here, we determined the cellular localization of Kcnj15 in the gills and examined its expression in response to changes in environmental salinity. In seawater-acclimated fish, Kcnj15 was localized to the basolateral membrane of Nka- and Nkcc1-positive ionocytes. Following acclimation to hypoosmotic brackish water (1 ppt), transcript levels of *cftr* and *kcnj1a* decreased, whereas *kcnj15* expression remained unchanged. Under this condition, Kcnj15 was localized to the basolateral membrane of Nka-positive ionocytes. These results demonstrate that Kcnj15 is localized to the basolateral membrane of gill ionocytes and suggest that it may function as a basolateral K^+^ leak channel supporting ionocyte homeostasis under both seawater and low-salinity conditions.

## 1. Introduction

The gill is a central osmoregulatory organ in teleost fishes and integrates ion transport with respiratory, acid-base, and nitrogen-excretory functions. In seawater (SW), teleosts maintain body fluid osmolarity at approximately one-third that of SW. To compensate for passive water loss and salt gain in the marine environment, they drink seawater and excrete excess ions through the gills, intestine, and kidneys (Evans et al., 2005; Marshall and Grosell, 2006; Takei, 2021). In the gills, mitochondrion-rich ionocytes secrete excess Na^+^ and Cl^−^ through a coordinated transport system in which basolateral Na^+^/K^+^-ATPase (Nka) establishes the Na^+^ gradient, basolateral Na^+^-K^+^-2Cl^−^ cotransporter 1 (Nkcc1) accumulates Cl^−^ within the cell, and apical cystic fibrosis transmembrane conductance regulator (Cftr) mediates Cl^−^ exit; Na^+^ is then secreted mainly through paracellular pathways (Hiroi et al., 2005; Inokuchi et al., 2008; Shaughnessy and Breves, 2025).

Transporting epithelial cells generally express high levels of Nka in the basolateral membrane, where it generates the electrochemical gradients that drive the secondary active transport of a wide variety of solutes. The basolateral membrane also expresses K^+^ channels that function as K^+^ leak channels (Enyedi and Czirják, 2010; Hebert et al., 2005; Heitzmann and Warth, 2008). These channels contribute to membrane potential generation by maintaining K^+^ permeability and support the continued operation of Nka by allowing K^+^ efflux from the cell. Nka and K^+^ leak channels are widely expressed not only in epithelial cells but also in nonepithelial cells, where they play fundamental roles in maintaining cellular homeostasis. In this respect, the basolateral membrane of transporting epithelia shares housekeeping functions with the plasma membrane of nonepithelial cells. Major classes of K^+^ leak channels include the Kcnk family, which encodes two-pore domain (K2P) K^+^ channels, and the Kcnj family, which encodes inwardly rectifying K^+^ (Kir) channels (Enyedi and Czirják, 2010; Silic et al., 2022). Members of the Kcnj family constitute one of the major classes of K^+^ channels involved in the maintenance of membrane potential and K^+^ homeostasis and are represented by 16 and 31 genes in humans and zebrafish, respectively (Manis et al., 2020; Silic et al., 2022). Expression of K^+^ channels in the basolateral membrane is a relatively common feature of transporting epithelia. In contrast, only a limited number of epithelial cell types express K^+^ channels in the apical membrane. Some transporting epithelial cells express K^+^ channels in both the apical and basolateral membranes, where they play important roles in transepithelial transport (Hebert et al., 2005; Heitzmann and Warth, 2008).

Various physiological roles of K^+^ channels have been described in fish. In gill ionocytes, both Nka and Nkcc1 transport K^+^ into the cell. Therefore, a conductive K^+^ exit pathway is required to stabilize intracellular K^+^, membrane potential, and the electrochemical driving forces for salt secretion. A classical electrophysiological study of the opercular epithelium of *Fundulus heteroclitus* showed that Ba^2+^-sensitive K^+^ conductance on the serosal side is coupled to Cl− secretion, providing early physiological evidence for such a basolateral K^+^-recycling pathway (Degnan, 1985). In the Japanese eel (*Anguilla japonica*), eKir (Kcnj16, Kir5.1), a hypertonicity-inducible inwardly rectifying K^+^ channel highly expressed in chloride cells, was identified as one of the first molecular candidates contributing to K^+^ conductance in ionocytes (Suzuki et al., 1999). In Mozambique tilapia (*Oreochromis mossambicus*), Kcnj1 (Kir1.1), also known as the renal outer-medullary potassium (Romk) channel, is localized to the apical membrane of gill ionocytes and mediates K^+^ excretion; expression of *kcnj1* paralogs is enhanced during exposure to elevated environmental K^+^ concentrations (Furukawa et al., 2014; Furukawa et al., 2012). A similar mechanism has been reported in Japanese medaka (*Oryzias latipes*), in which ionocytes actively secrete K^+^ and acclimation to high-K^+^ conditions increases branchial expression of *kcnj1a* (*romka*) and *nkcc1a* (Horng et al., 2017). More recently, salinity-dependent regulation of *kcnj1a* expression has also been reported in Asian sea bass (*Lates calcarifer*) (Ding et al., 2025).

Additional K^+^-channel families have also been implicated in osmoregulatory epithelia. In Atlantic salmon *Salmo salar*, gill BK-channel transcripts respond dynamically to transfer from freshwater (FW) to brackish water (BW), with a transient increase after osmotic challenge (Loncoman et al., 2018).

Single-nucleus transcriptomic data from Atlantic salmon gills further identified Kir4.2/Kcnj15 as one of the prominent K^+^ -channel transcripts in SW-type ionocytes, although that study did not determine the cellular localization of the Kcnj15 protein (Shaughnessy and Breves, 2025; West et al., 2021). In the shark rectal gland, a specialized salt-secreting epithelium, the basolateral two-pore-domain channel TASK-1 provides a K^+^ conductance required to sustain Cftr-dependent Cl^−^ secretion (Telles et al., 2016). These observations indicate that K^+^ channels supporting salt-transporting epithelia have been recruited from multiple channel families and that their membrane distribution and regulation are species dependent.

The diversity of fish inwardly rectifying K^+^ (Kir/Kcnj) channels extends beyond osmoregulatory tissues. Zebrafish *Danio rerio* possess an expanded Kir repertoire with distinct developmental expression domains (Abbas et al., 2011; Silic et al., 2022). In the heart of rainbow trout (*Oncorhynchus mykiss*), Kcnj2.1 (Kir2.1) and Kcnj2.2 (Kir2.2) expression changes with thermal acclimation (Hassinen et al., 2007), whereas the novel Kcnj17 (Kir2.5) channel of crucian carp (*Carassius carassius*) is upregulated during chronic cold exposure (Hassinen et al., 2008). Together with the branchial studies described above, these findings illustrate extensive paralog-, tissue-, and species-specific use of K^+^ channels in fishes and argue against assuming that a single mammalian K^+^ -channel model applies across teleost osmoregulatory epithelia.

In Japanese pufferfish *Takifugu rubripes*, our previous study identified Kcnj15/Kir4.2 as a candidate K^+^ channel involved in epithelial ion transport in the kidney, urinary bladder, and intestine (Sakamoto et al., 2026). Interestingly, Kcnj15 was localized to either the apical and/or basolateral membrane of the epithelial cells, and its localization varied depending on the tissue type and environmental salinity. Thus, whether Kcnj15 is expressed in ionocytes, which membrane domain it occupies, and whether its branchial expression changes with environmental salinity remained unknown.

Japanese pufferfish are euryhaline and tolerate markedly diluted seawater (Lee et al., 2005). Here, fish were acclimated to 1-ppt brackish water to examine branchial responses to a low-salinity environment. We quantified gill mRNA levels of *kcnj1a, kcnj1b, kcnj15, nkcc1a, nkcc1b, nkcc2, ncc*, and *cftr* in pufferfish acclimated to SW, KCl-supplemented SW, 1-ppt brackish water, or KCl-supplemented 1-ppt brackish water. We further examined Kcnj15 immunolocalization relative to anti-Nkcc (T4) and Nka signals in fish acclimated to SW or 1-ppt brackish water. These analyses revealed that Kcnj15 is localized to the basolateral membrane of gill ionocytes in the Japanese pufferfish and suggest that it may represent one of the major basolateral K^+^ leak channels in these cells.

## 2. Materials and methods

### 2.1. Animals

Japanese pufferfish (*Takifugu rubripes*) were purchased from a local dealer. For real-time PCR experiments, Japanese pufferfish (24–41 g; n = 4 for each condition) were reared in 150-L tanks containing natural SW (35 ppt), natural SW supplemented with 10 mM KCl (SW-HK; Nacalai, Kyoto, Japan), 1-ppt brackish water (BW) prepared by diluting natural SW approximately 35-fold with dechlorinated tap water, or 1-ppt BW supplemented with 2 mM KCl (BW-HK) for 7–11 d as previously described (Sakamoto et al., 2026). For immunohistochemical experiments, Japanese pufferfish (300–400 g) were reared in 150-L tanks containing natural SW (35 ppt) or 1-ppt BW for 7–11 d as previously described (Sakamoto et al., 2026). The pufferfish were fed once daily and starved for 2 d before the experiments. All experimental animals were anesthetized via immersion in 0.1% ethyl m-aminobenzoate (MS-222; tricaine; Sigma-Aldrich, St. Louis, MO, USA) neutralized with sodium bicarbonate. After humane killing via cervical transection, the gills were dissected. All pufferfish were housed and cared for in accordance with the manual approved by the Institutional Animal Experiment Committee of the Institute of Science Tokyo.

### 2.2. Antibodies

Polyclonal antiserum against the Japanese pufferfish Kcnj15 was prepared using rabbits immunized with bovine serum albumin-conjugated synthetic peptides corresponding to the 14 amino acid carboxyl terminus of Japanese pufferfish Kcnj15 (Sakamoto et al., 2026). The anti-Nkcc mouse monoclonal antibody (clone T4) was purchased from Sigma-Aldrich (P/N MABS1237). The anti-Nkcc (T4) antibody recognizes Nkcc1, Nkcc2, and Ncc (Hiroi et al., 2008; Lytle et al., 1995). Anti-eel Na^+^ /K^+^ -ATPase (Nka) rat antiserum was obtained from previous studies (Kato et al., 2011; Miyamoto et al., 2002). Preimmune serum collected from the same rabbit was used as the negative control for the anti-Kcnj15 antibody, and normal mouse or rat serum was used as the negative control for the anti-Nkcc (T4) or anti-Nka antibody, respectively. The antigen-purified anti-Kcnj15 antibody and antigen-absorbed anti-Kcnj15 antiserum was prepared as previously described (Sakamoto et al., 2026).

### 2.3. Immunohistochemistry

The gills of Japanese pufferfish were perfused and fixed with 4% paraformaldehyde (PFA) in 100 mM phosphate buffer (pH 7.4) and then excised as previously described (Kato et al., 2007; Mistry et al., 2004). The tissues were further fixed in the same fixative at 4°C for 2 h, rinsed in phosphate-buffered saline (PBS), cryoprotected in a graded series of sucrose solutions up to 20%, and rapidly frozen in optimal cutting temperature (OCT) compound (Sakura Finetek, Tokyo, Japan).

Immunohistochemical analyses were performed on frozen gill sections (6 μm) obtained from Japanese pufferfish acclimated to SW or 1-ppt BW as previously described (Sakamoto et al., 2026). The sections were treated with phosphate-buffered saline (PBS) containing 0.1% Triton X-100 for 10 min at 23 °C and blocked with PBS containing 5% fetal bovine serum (FBS) for 1 h at 23 °C. For evaluation of anti-Kcnj15 staining specificity, sections were incubated with anti-Kcnj15 rabbit serum (1:1,000 dilution), antigen-purified anti-Kcnj15 antibody (0.46 μg/mL), antigen-absorbed anti-Kcnj15 antiserum (1:1,000 dilution), or preimmune rabbit serum (1:1,000 dilution) as previously described (Sakamoto et al., 2026). For double immunostaining, sections were incubated with the antigen-purified anti-Kcnj15 antibody (0.46 μg/mL) together with the anti-Nkcc (T4) mouse monoclonal antibody (0.8 μg/mL) or anti-Na^+^ /K^+^ -ATPase (Nka) rat antiserum (1:1,000 dilution). Normal mouse or rat serum (1:1,000 dilution) was used as a negative control for the anti-Nkcc (T4) or anti-Nka antibody, respectively. All primary antibody incubations were performed for 16 h at 23 °C. After washing with PBS, sections were stained with Alexa Fluor 488-labeled F(ab’)_2_ fragment of donkey anti-rabbit IgG, Cy3-labeled F(ab’)_2_ fragment of goat anti-mouse IgG, or Cy3-labeled F(ab’)_2_ fragment of goat anti-rat IgG (each 1:2,000; Jackson ImmunoResearch Laboratories, West Grove, PA, USA), and Hoechst 33342 (100 ng/mL). Fluorescence images were obtained using a confocal laser scanning microscope (CLSM, LSM780; Carl Zeiss, Oberkochen, Germany) and processed using ZEN software (Carl Zeiss). Images of the test samples and corresponding negative controls were acquired under identical detector gain and laser power settings. The original images for each fluorescence channel were acquired in grayscale and are displayed in pseudo-color.

### 2.4. Real-time PCR

Total RNA was extracted from the gills using Isogen (Nippon Gene, Tokyo, Japan), and first-strand cDNA was synthesized from 5 μg of total RNA using the SuperScript IV First-Strand Synthesis System (Thermo Fisher Scientific, Waltham, MA, USA) with oligo(dT) primers as previously described (Sakamoto et al., 2026). The relative expressions of *kcnj1a, kcnj1b, kcnj15, nkcc1a, nkcc1b, nkcc2, ncc*, and *cftr* were quantified by real-time PCR using the SYBR Green method on a Thermal Cycler Dice Real Time System III (Takara Bio, Shiga, Japan), with *actb* as the reference gene and gene-specific primers (Table 1). Significant differences at *p* < 0.05 were determined separately for each gene using one-way analysis of variance (ANOVA), followed by the Tukey–Kramer multiple comparisons test (GraphPad Prism 5, GraphPad Software, San Diego, CA, USA). Reproducibility was confirmed using two sets of experiments.

**Table 1.**
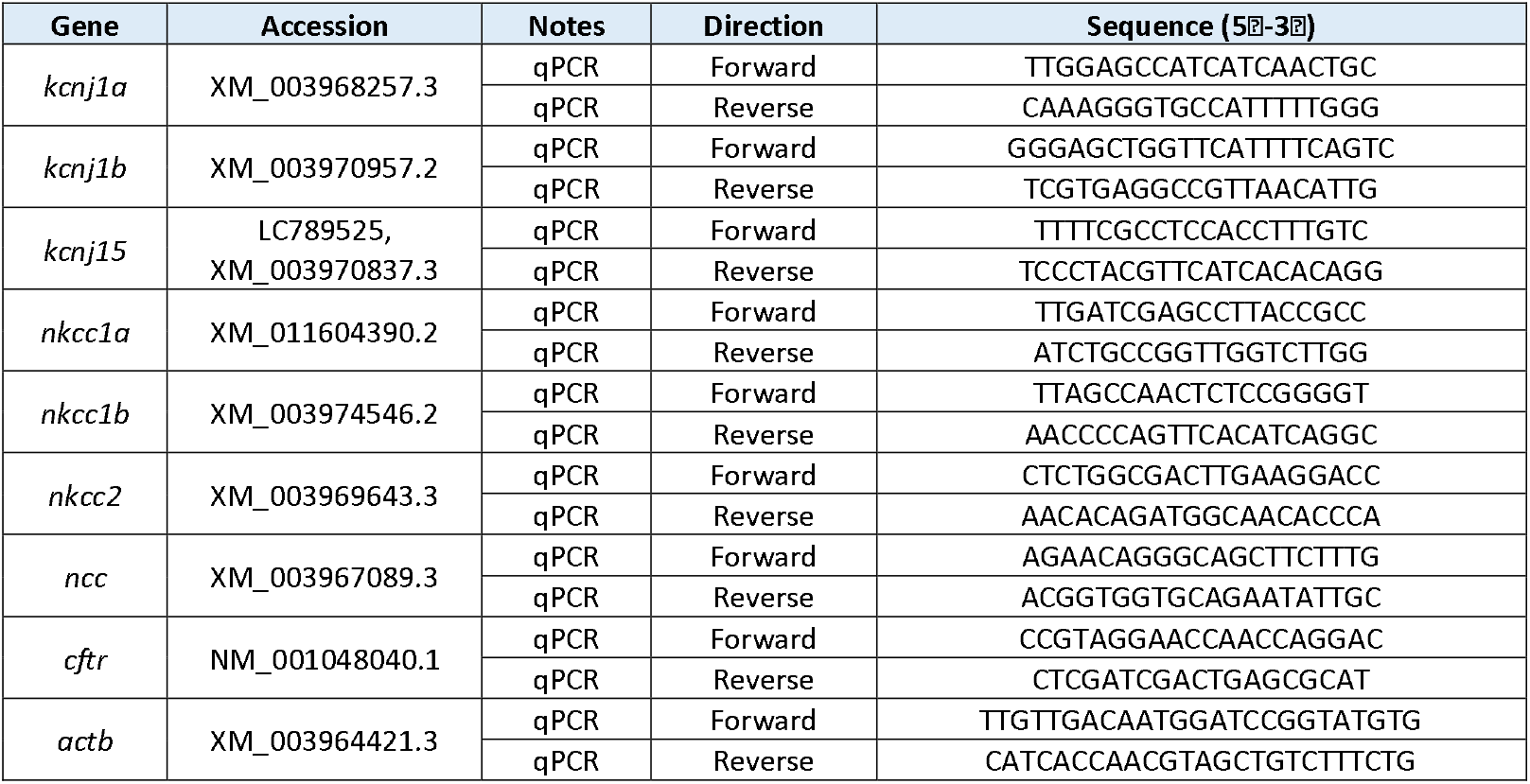
Primers used for real-time PCR in this study.

## 3. Results

### 3.1. Expression Kcnj15 and other ion transporters in the gills of Japanese pufferfish

Previous semiquantitative RT-PCR analyses using various tissues of the Japanese pufferfish showed that *kcnj15* is expressed in the gills (Sakamoto et al., 2026). To gain insight into the function of *kcnj15* in the gills, we quantified *kcnj15* expression by quantitative PCR in fish acclimated to seawater (SW), SW supplemented with 10 mM KCl (SW-HK), 1-ppt brackish water (BW), or 1-ppt BW supplemented with 2 mM KCl (BW-HK) (Fig. 1). Expression levels of *kcnj1a, kcnj1b, nkcc1a, nkcc1b, nkcc2, ncc*, and *cftr* were analyzed in parallel.

**Figure 1.**
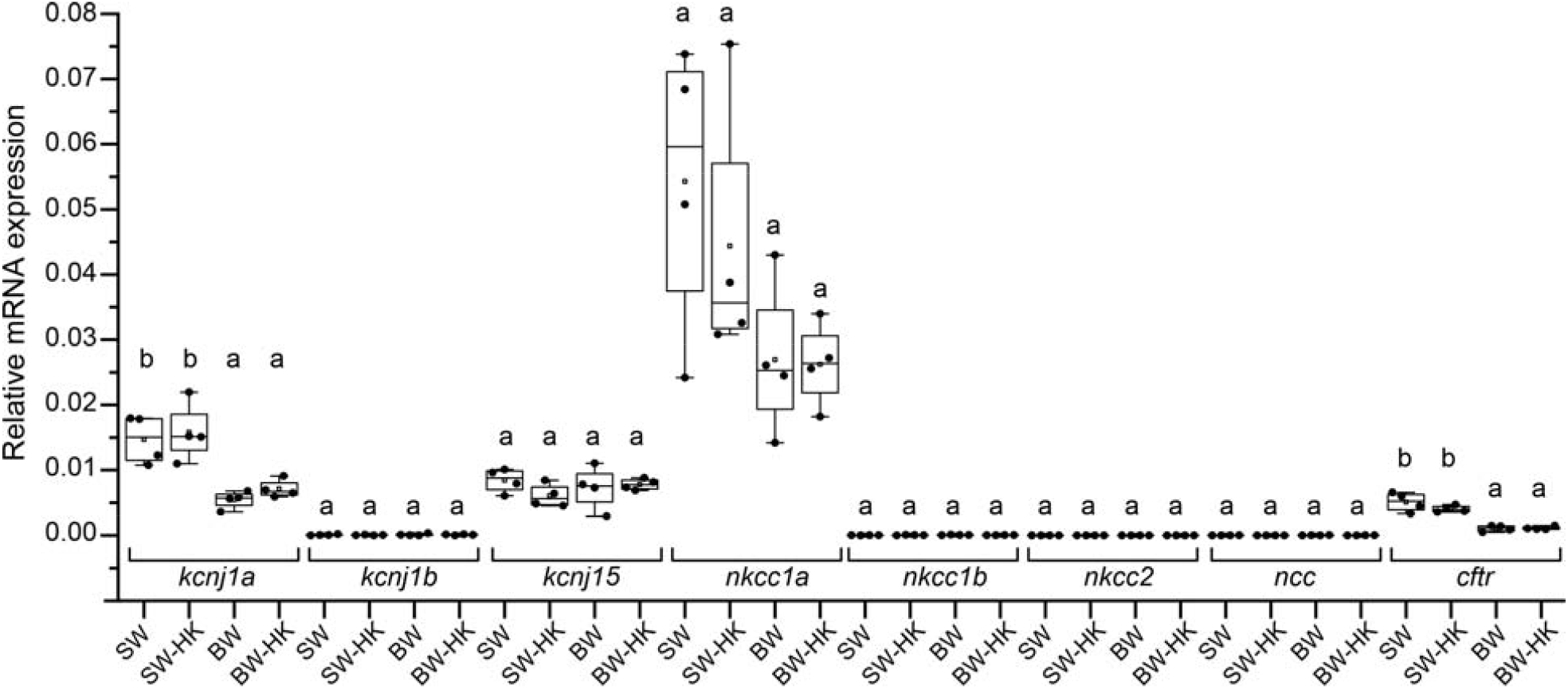
Real-time PCR quantification of mRNAs for *kcnj1a, kcnj1b, kcnj15, nkcc1a, nkcc1b, nkcc2, ncc*, and *cftr* in the gills of Japanese pufferfish acclimated to SW, SW supplemented with 10 mM KCl (SW-HK), 1-ppt BW, or 1-ppt BW supplemented with 2 mM KCl (BW-HK). Values are expressed relative to *actb* and presented as box-and-whisker plots (*n* = 4 per group). Black dots indicate individual values, open squares indicate means, boxes indicate interquartile ranges, horizontal lines within boxes indicate medians, and whiskers indicate minimum and maximum values. Significant differences at *p* < 0.05 were determined separately for each gene using one-way ANOVA followed by the Tukey–Kramer multiple comparisons test. Groups labeled with different letters indicate statistically significant differences.

*kcnj15* mRNA was detected under all conditions, with no significant difference among the SW, SW-HK, 1-ppt BW, and BW-HK groups (Fig. 1). In contrast, mRNA levels of another K^+^ channel gene, *kcnj1a*, were significantly higher under SW and SW-HK conditions than under 1-ppt BW and BW-HK conditions. KCl supplementation did not produce a clear change within the same salinity condition. *kcnj1b* mRNA levels were very low under all conditions and did not differ significantly among the four groups.

*cftr* mRNA levels were significantly higher under SW and SW-HK conditions than under 1-ppt BW and BW-HK conditions. No clear effect of KCl supplementation was observed within the same salinity condition. *nkcc1a* was expressed at a relatively high level among the ion transport-related genes examined, but its expression did not differ significantly among the four groups. In contrast, *nkcc1b, nkcc2*, and *ncc* mRNA levels were very low under all conditions (Fig. 1).

These analyses revealed that *kcnj1a, cftr, kcnj15*, and *nkcc1a* are highly expressed in the gills of the Japanese pufferfish. In addition, expression of *kcnj1a* and *cftr* was reduced under low-salinity conditions, whereas expression of *kcnj15* and *nkcc1a* remained largely unchanged despite changes in environmental salinity.

### 3.2. Basolateral localization of Kcnj15 in gill ionocytes of SW-acclimated Japanese pufferfish

To determine which cell types express Kcnj15 in the gills and to examine its subcellular localization, immunohistochemical staining was performed on gill sections of the Japanese pufferfish using an antibody against pufferfish Kcnj15. The specificity of Kcnj15 immunostaining in the gill was evaluated using sections from SW-acclimated fish. Kcnj15 immunosignals were detected in gill epithelial cells with both the antigen-purified anti-Kcnj15 antibody and anti-Kcnj15 antiserum. In contrast, no comparable signals were observed with the antigen-absorbed anti-Kcnj15 antiserum or preimmune serum (Fig. 2A).

**Figure 2.**
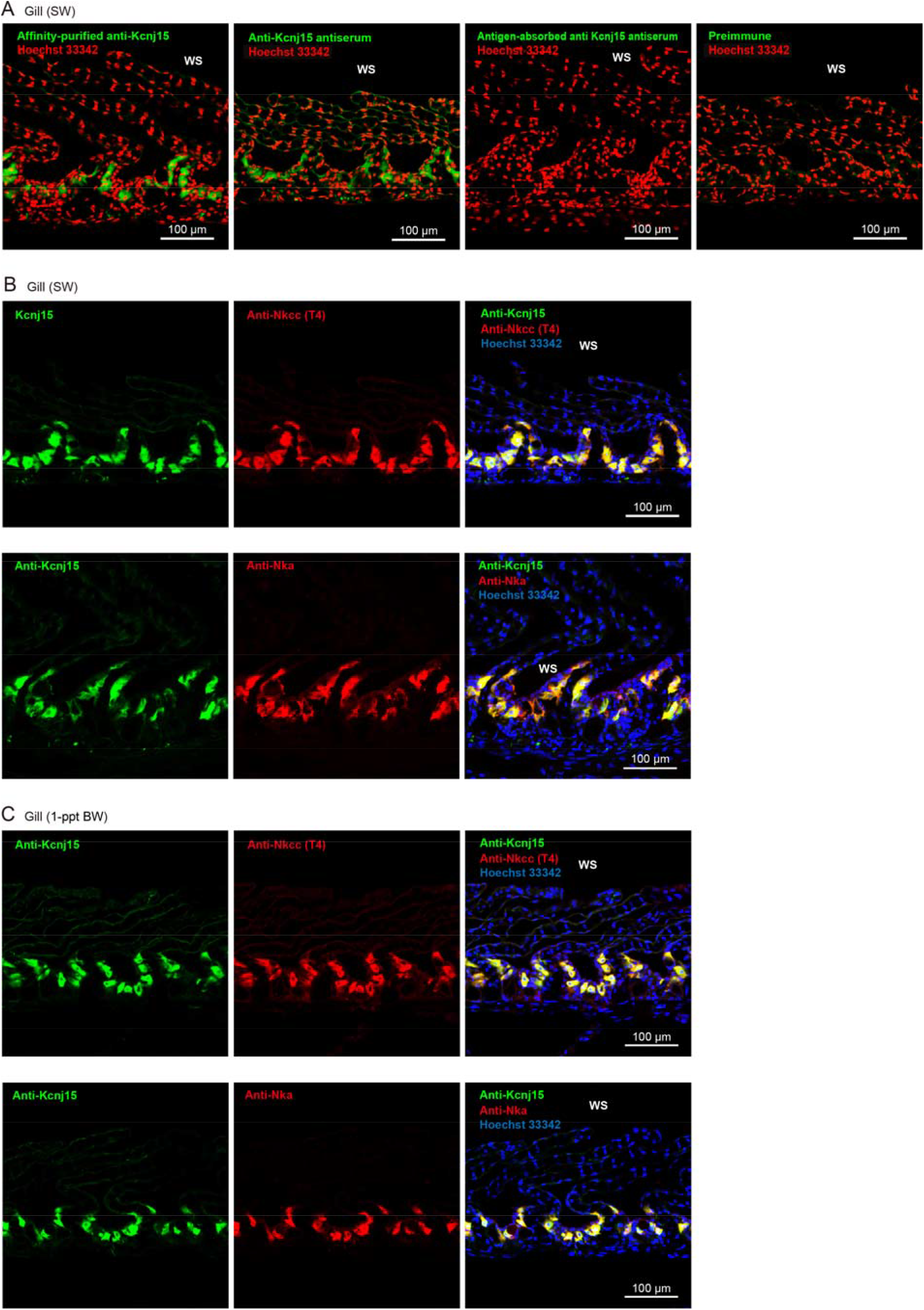
Localization of Kcnj15 in the gills of SW- and 1-ppt BW-acclimated Japanese pufferfish. (A) Immunohistochemical analysis of SW-acclimated gill sections with the antigen-purified anti-Kcnj15 antibody, anti-Kcnj15 antiserum, antigen-absorbed anti-Kcnj15 antiserum, and preimmune serum. Kcnj15 immunosignals are shown in green, and nuclei stained with Hoechst 33342 are shown in red. (B, C) Gill sections from SW-acclimated (B) and 1-ppt BW-acclimated (C) pufferfish were stained with the antigen-purified anti-Kcnj15 antibody (green) and anti-Nkcc (T4) (red) or anti-Na^+^ /K^+^ -ATPase (Nka) (red) antibodies. Nuclei were stained with Hoechst 33342 (blue) in merged images. The anti-Nkcc (T4) antibody recognizes Nkcc1, Nkcc2, and Ncc; because *nkcc1a* was expressed at a relatively high level in the gill, whereas *nkcc1b, nkcc2*, and *ncc* showed very low expression, the anti-Nkcc (T4) signal in the gill is interpreted to predominantly reflect Nkcc1a. Scale bars: 100 μm. WS, water side.

In SW-acclimated fish, Kcnj15 signals were observed in a subset of gill epithelial cells and showed substantial overlap with signals detected by the anti-Nkcc (T4) antibody (Fig. 2B). Because the anti-Nkcc (T4) antibody recognizes Nkcc1, Nkcc2, and Ncc, and because *nkcc1a* was expressed at a relatively high level in the gill whereas *nkcc1b, nkcc2*, and *ncc* were expressed at very low levels, the anti-Nkcc (T4) signal in the gill was interpreted as predominantly reflecting Nkcc1a. Kcnj15 signals also substantially overlapped with Na^+^ /K^+^ -ATPase (Nka)-positive areas (Fig. 2B). These staining patterns indicate that Kcnj15 is localized to the basolateral membrane of gill ionocytes in SW-acclimated Japanese pufferfish.

### 3.3. Basolateral localization of Kcnj15 in gill ionocytes of Japanese pufferfish under low-salinity condition

Kcnj15 localization was similarly examined in the gills of pufferfish acclimated to a hypoosmotic environment. A similar distribution was observed in fish acclimated to 1-ppt BW. Kcnj15 signals were detected in a subset of gill epithelial cells and substantially overlapped with both anti-Nkcc (T4) and Nka signals (Fig. 2C). These findings indicate that Kcnj15 is also localized to the basolateral membrane of gill ionocytes under 1-ppt BW conditions, with no apparent salinity-dependent change in its cellular distribution compared with that in SW-acclimated fish.

## 4. Discussion

K^+^ channels are involved in a variety of epithelial transport processes underlying osmoregulation in fish. Although numerous studies, including those described in the Introduction, have investigated the physiological roles of K^+^ channels in fish, many aspects of their functions and regulation remain poorly understood. In particular, Kcnj15 has received little attention in teleost fishes. Recently, we identified Kcnj15 as a K^+^ channel expressed in several osmoregulatory organs of the Japanese pufferfish, including the gills, intestine, kidney, and urinary bladder, and characterized its cellular localization in the intestine, kidney, and urinary bladder (Sakamoto et al., 2026). Interestingly, Kcnj15 was localized to both the apical and basolateral membranes of intestinal epithelial cells and collecting duct cells in SW-acclimated fish, whereas it was restricted to the basolateral membrane in the collecting ducts and urinary bladder of fish acclimated to hypoosmotic conditions (Sakamoto et al., 2026). These findings indicated that the subcellular localization of Kcnj15 varies depending on both cell type and environmental salinity and suggest that Kcnj15 is one of the key factors involved in osmoregulation in teleost fish. Previous studies on branchial K^+^ channels in teleosts reported the expression of Kcnj16 in the basolateral membrane (Suzuki et al., 1999) and Kcnj1 in the apical membrane of ionocytes (Furukawa et al., 2014; Furukawa et al., 2012). However, the cellular localization of Kcnj15 in fish gills had not been examined. In the present study, immunohistochemical analyses using gill sections and a specific anti-pufferfish Kcnj15 antibody demonstrated that Kcnj15 is localized to the basolateral membrane of branchial ionocytes. Furthermore, neither the expression level nor the cellular localization of Kcnj15 was affected by environmental salinity. These findings provide new insight into the molecular organization of branchial ionocytes and identify Kcnj15 as a previously unrecognized component of the basolateral transport machinery in teleost gills.

The high expression of *cftr, kcnj1a*, and *nkcc1a* and the localization of Nkcc1a and Nka indicate that Japanese pufferfish possess the canonical SW-type ionocyte transport machinery described in other marine teleosts (Evans et al., 2005; Hiroi and McCormick, 2012; Shaughnessy and Breves, 2025). This mechanism shares many features with the NaCl secretory mechanism of the rectal gland in cartilaginous fishes (Silva and Evans, 2024; Telles et al., 2016). Moreover, the higher expression of *cftr* and *kcnj1a* under SW conditions than under hypoosmotic conditions is consistent with their proposed roles in branchial salt secretion. Together, these results indicate that the salt-secretory machinery of Japanese pufferfish ionocytes is broadly similar to that of other marine teleosts (Fig. 3A). In addition, the basolateral localization of Kcnj15 identified in the present study suggests that Kcnj15 may function as one of the K^+^ leak channels that support sustained Nka activity and maintenance of membrane potential. Thus, Kcnj15 is likely to represent a previously unrecognized component of the ion-secretion machinery in SW-type ionocytes.

**Figure 3.**
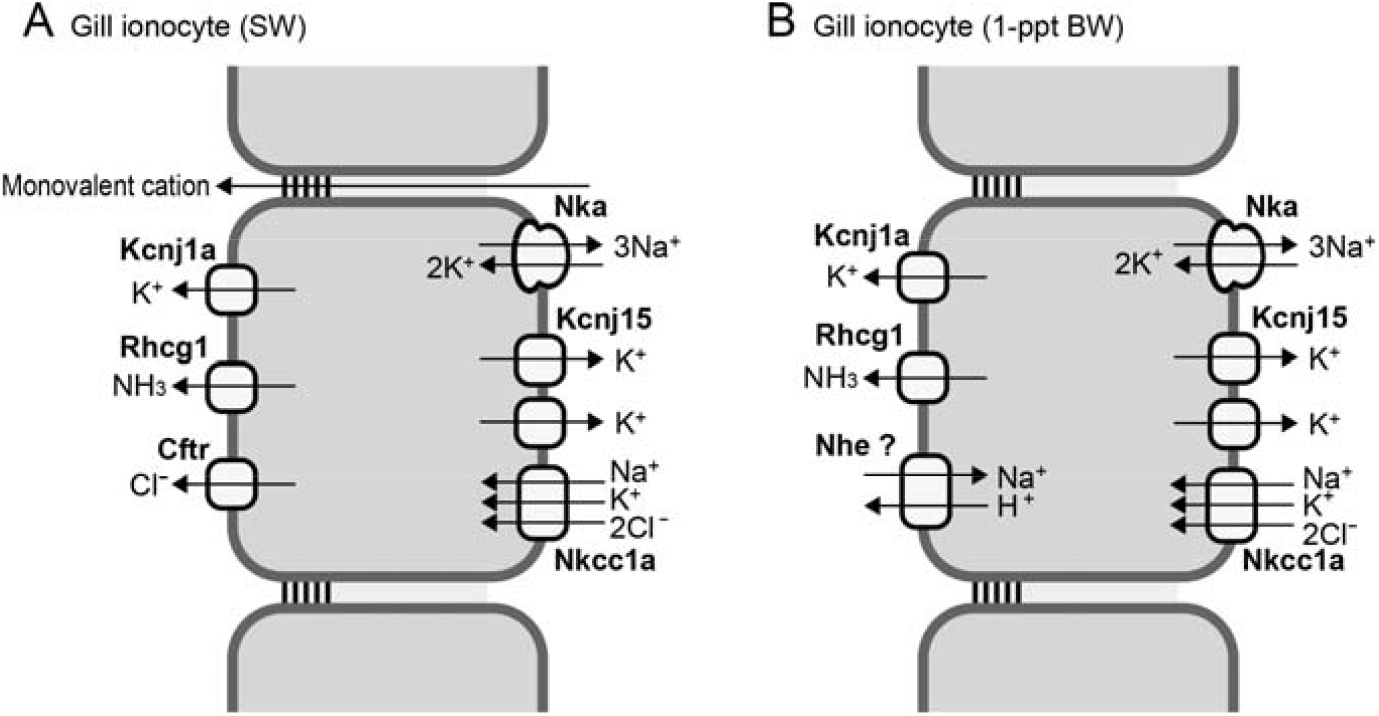
Proposed model of Kcnj15 as one of the basolateral K^+^ leak channels in pufferfish ionocytes. The localization of Kcnj15, Nka, and Nkcc1a was shown in this study. (A) Model depicting the proposed role of Kcnj15 in gill ionocytes of SW-acclimated Japanese pufferfish. Rhcg1 is positioned based on previous studies of Japanese pufferfish (Nakada et al., 2007) and other teleosts (Shih et al., 2013; Shih et al., 2012), and Kcnj1a and Cftr are positioned based on previous studies of other teleosts (Furukawa et al., 2014; Furukawa et al., 2012; Hiroi et al., 2005). (B) Model depicting the proposed role in gill ionocytes of 1-ppt BW-acclimated Japanese pufferfish. Rhcg1 and Kcnj1a are positioned based on previous studies of other freshwater teleosts (Furukawa et al., 2014; Furukawa et al., 2012; Horng et al., 2017; Shih et al., 2013; Shih et al., 2012). The mechanism of NaCl uptake in ionocytes of 1-ppt BW-acclimated Japanese pufferfish remains unclear. Because the Japanese pufferfish lacks the *ncc2* gene, Na^+^/H^+^ exchanger (e.g. Nhe3) is thought to be the major Na^+^ uptake pathway, as in other freshwater teleosts (Inokuchi et al., 2009; Inokuchi et al., 2008; Inokuchi et al., 2017; Shih et al., 2012); however, this has not been fully investigated.

The Japanese pufferfish is a marine teleost, but it can survive in highly diluted SW under hypoosmotic conditions (Lee et al., 2005). When acclimated to a hypoosmotic environment, Japanese pufferfish produce hypotonic urine (Sakamoto et al., 2026), as reported for many freshwater (FW) teleosts, indicating that the kidney adopts a FW-like osmoregulatory function under such conditions. In contrast, little is known about the characteristics of branchial ionocytes in hypoosmotic environments, and it remains unclear whether Japanese pufferfish develop FW-type ionocytes comparable to those described in other FW teleosts. Previous studies have revealed substantial diversity in FW-type ionocytes among teleost species, including differences in ionocyte subtypes and their distribution patterns (Hiroi and McCormick, 2012; Inokuchi et al., 2009; Inokuchi et al., 2017). Two major FW-type ionocyte types are characterized by apical expression of either Na^+^/H^+^ exchanger 3 (Nhe3) or Na^+^-Cl− cotransporter 2 (Ncc2). Because the Japanese pufferfish lacks the *ncc2* gene (Motoshima et al., 2023; Ota et al., 2024), an Ncc2-dependent ion uptake pathway is unlikely to be present in this species. To gain insight into the function of branchial Kcnj15, we analyzed gills from fish acclimated to a hypoosmotic environment. Immunohistochemical analyses demonstrated that Kcnj15, Nka, and Nkcc1a remained localized to the basolateral membrane of branchial ionocytes. These observations suggest that, even under hypoosmotic conditions, Kcnj15 may function as a basolateral K^+^ leak channel that supports sustained Nka activity and membrane potential maintenance. Interestingly, Nkcc1a, which is generally regarded as a basolateral Cl− uptake pathway for salt secretion in seawater-type ionocytes, also remained localized to the basolateral membrane and maintained its mRNA expression under hypoosmotic conditions. These findings suggest that basolateral Nkcc1a retains physiological functions in ionocytes even in low-salinity environments. One possible role of Nkcc1a under these conditions may be related to ammonia transport. In Nhe3-expressing FW-type ionocytes, the ammonia channel Rhcg1 and Nhe3 facilitate NH_3_/NH_4_^+^ excretion coupled to Na^+^ uptake and thereby contributes to ion acquisition from dilute environments (Ito et al., 2014; Kumai and Perry, 2011; Nakada et al., 2007; Shih et al., 2013; Shih et al., 2012). Basolateral NH_4_^+^ entry has been proposed to occur through transport systems that can substitute NH_4_^+^ for K^+^, including K^+^ channels and Nka. Because Nkcc1a is also a K^+^ transporter, it could potentially contribute to NH_4_^+^ uptake across the basolateral membrane (Bergeron et al., 2003). However, transporters localized to the apical membrane of ionocytes in hypoosmotic-acclimated pufferfish have not yet been characterized, and their identification will be an important subject for future studies.

A useful comparison is provided by the closely related river puffer *Takifugu obscurus*, which can inhabit both SW and FW environments (Kato et al., 2005). In this species, transfer from FW to SW increases branchial *nkcc1* and *nka* expression and is accompanied by a shift from FW-associated lamellar ionocytes to seawater-associated filament ionocytes (Ding et al., 2020). In contrast, only filament ionocytes were observed in the gills of Japanese pufferfish under both seawater and hypoosmotic conditions (Fig. 2). These differences suggest that the mechanisms of hypoosmotic adaptation differ between the marine Japanese pufferfish, which cannot survive in pure FW, and the more euryhaline river puffer, which tolerates a much wider salinity range.

KCl supplementation produced no clear expression change within the same salinity condition for the genes examined. This result is consistent with our previous study on the pufferfish kidney, intestine, and urinary bladder, which also found no significant transcriptional response to the K^+^ supplementation used in these experiments (Sakamoto et al., 2026). Previous studies in tilapia (Furukawa et al., 2014; Furukawa et al., 2012) and medaka (Horng et al., 2017) have demonstrated transcriptional responses of *kcnj1* to elevated environmental K^+^ concentration. Our KCl additions were intentionally modest, and K^+^ concentration, exposure duration, sampling time, and species-specific regulatory thresholds could all influence the transcriptional response. Preliminary observations suggest that Japanese pufferfish have limited tolerance to changes in environmental K^+^ concentration, making acclimation experiments under a wide range of K^+^ conditions difficult to conduct (Sakamoto et al., 2026). The data obtained in the present study are insufficient to draw firm conclusions regarding the roles of Kcnj15 and other membrane transporters in responses to altered environmental K^+^ levels. Nevertheless, future studies addressing this question will require careful consideration and optimization of experimental conditions.

## 5. Conclusion

In conclusion, the present study identifies Kcnj15 as a K^+^ -channel protein associated with the basolateral region of Nka/Nkcc1a-positive gill ionocytes in Japanese pufferfish and shows that this distribution is retained after acclimation from SW to 1-ppt brackish water. In parallel, *kcnj1a* and *cftr* expression decreased at low salinity, whereas *kcnj15* and *nkcc1a* expression did not differ significantly among salinity and KCl conditions. We propose that pufferfish Kcnj15 may function as a basolateral K^+^ leak pathway that supports ionocyte homeostasis under both SW and low-salinity conditions.

## CRediT authorship contribution statement

**Yohei Sakamoto:** Conceptualization, Data curation, Formal analysis, Investigation, Methodology, Project administration, Visualization, Writing – original draft, Writing – review & editing. *Akira Kato:* Conceptualization, Funding acquisition, Methodology, Supervision, Writing – original draft, Writing – review & editing.

## Ethics declaration

All pufferfish were housed and cared for in accordance with the protocol approved by the Institutional Animal Experiment Committee of the Institute of Science Tokyo.

## Acknowledgements

We would like to thank Dr. Ayumi Nagashima, the Bioscience Center, and the Open Research Facilities for Life Science and Technology at the Institute of Science Tokyo for their technical assistance in this study.

## Data availability

Data will be made available on request.

